# SweHLA: the high confidence HLA typing bio-resource drawn from 1 000 Swedish genomes

**DOI:** 10.1101/660241

**Authors:** Jessika Nordin, Adam Ameur, Kerstin Lindblad-Toh, Ulf Gyllensten, Jennifer R.S. Meadows

**Author notes:** Corresponding author: Jessika Nordin. **Funding**: This study was supported by Knut and Alice Wallenberg Foundation (2018.0101) and Vetenskapsrådet Rådsprofessor (541-2013-8161). The funders had no role in study design, data collection and analysis, decision to publish, or preparation of the manuscript.

## Abstract

There is a need to accurately call human leukocyte antigen (HLA) genes from existing short-read sequencing data, however there is no single solution that matches the gold standard of lab typing. Here we aimed to combine results from available software, minimising the biases of applied algorithm and HLA reference. The result is a robust HLA population resource for the published 1 000 Swedish genomes, and a framework for future HLA interrogation. HLA 2-field alleles were called using four imputation and inference methods for the classical eight genes (class I: *HLA-A, -B, -C*; class II: *HLA-DPA1, -DPB1, -DQA1, -DQB1, -DRB1*). A high confidence population set (SweHLA) was determined using an *n-1* concordance rule for class I (four software) and class II (three software) alleles. Results were compared across populations and individual programs benchmarked to SweHLA. Per allele, 875 to 988 of the 1 000 samples were genotyped in SweHLA; 920 samples had at least seven loci. While a small fraction of reference alleles were common to all software (class I=1.9% and class II=4.1%), this did not affect the overall call rate. Gene-level concordance was high compared to European populations (>0.83%), with COX and PGF the dominant SweHLA haplotypes. We noted that 15/18 discordant alleles (delta allele frequency > 2) were previously reported as disease-associated. These differences could in part explain across-study genetic replication failures, reinforcing the need to use multiple software. SweHLA demonstrates a way to use existing NGS data to generate a population resource agnostic to individual HLA software biases.

## Introduction

The human major histocompatibility complex (MHC) spans approximately four Mb on chromosome six and contains more than 200 genes, ∼40% of which have immunological function ^1^. Within this region, the human leukocyte antigen (HLA) genes are divided into classes (I, II and III). These are some of the most polymorphic genes in the genome, with new alleles continuously being discovered ^1–3^. In the ten years from 2008 to 2018, the number of alleles in class I and class II have expanded six fold; from ∼2 500 to ∼15 500, and ∼1 000 to ∼6 000 alleles, respectively ^4^. Given their roles in immune recognition (intracellular, class I, and extracellular, class II), these genes are essential to the processes of transplantation, disease and infection susceptibility (including immunological diseases, but also cancers and neuropathies), drug response and pregnancy ^2,3,5,6^.

Lab typing is the gold standard for calling HLA alleles, where alleles are usually called at the clinically relevant, protein level, 2-field resolution ^7^. However, the last ten years has seen the growing need to accurately call alleles from pre-existing data, such as that generated from SNP chips or NGS short-read sequencing ^7^. The result has been an explosion of HLA software solutions, each using different methods for imputation or inference. The continued growth in this bioinformatics field neatly illustrates the difficulty of the task, and demonstrates how, as yet, no single software can replace biological typing.

Using four freely available software, and existing Illumina short read NGS data generated for the 1 000 Swedish genomes project (SweGen ^8^), we called 2-field alleles for the classical eight HLA genes (class I: *HLA-A, -B, -C*; class II: *HLA-DPA1, -DPB1, -DQA1, -DQB1, -DRB1*). This multi-software dataset demonstrated how biases inherent in input data choice, HLA allele reference availability and software algorithms, could impact downstream analyses. For these reasons, alleles within the high confidence Swedish population HLA set, SweHLA, were designated on the basis of *n-1* software matches (class I: three out of four; class II: two out of three). This resource, benchmarked with allele frequency correlation to 245 previously lab typed Swedish individuals ^9,10^ and compared on a population level to 5 544 imputed British individuals ^11^, is publicly available for research use.

## Methods

### Study population

Individual bam and gVCF files from SweGen ^8^ were used as a basis for the analyses. Representing a cross-section of the Swedish population, these individuals were selected from the Swedish twin registry (one per pair) and The Northern Sweden Population Health Study (in total 506 males and 494 females with a median age of 65.2 years) ^8^. Illumina HiSeq X data had an average genome coverage of 36.7 x ^8^.

### MHC demographics

The MHC region was defined as spanning hg19 chr6:28 477 797-33 448 354 using coordinates lifted from GRCh38.p13. Nucleotide diversity (Pi), Tajima’s D, and SNP and indel densities were calculated in 1 000 bp windows from curated vcfs using VCFtools ^12^ v0.1.14. Coverage across the same windows was determined with BEDtools ^13^ v2.26.0 using individual sorted bam files and a read length of 150 bp ^8^.

### HLA typing with four software

Four freely available software programs were selected for the analysis (Figure 1); the commonly used imputation (SNP2HLA ^14^, cited >340 times) and inference software (OptiType ^15^, cited >140 times), as well as two more recently published inference solutions (HLA-VBSeq ^16^ and HLAscan ^17^). In brief, the imputation method builds HLA alleles based on haplotypes generated from user supplied pruned GWAS SNPs and a phased reference panel. Whereas inference software aligns NGS reads to all HLA alleles in a reference and determines an allele best match via method specific penalty algorithms. The reference is sourced from the ImMunoGeneTics project/human leukocyte antigen (IMGT/HLA) database ^18^. Of note, each software method uses a different reference version, and different regions of this resource, be it nucleotide (exonic) or genomic sequence. The 2-field resolution alleles from each program were recorded for each HLA gene available, however only alleles from the classical 8 genes were evaluated for the generation of SweHLA. The specific running conditions of each software is detailed below and summarised in Figure 1.

**Figure 1.**
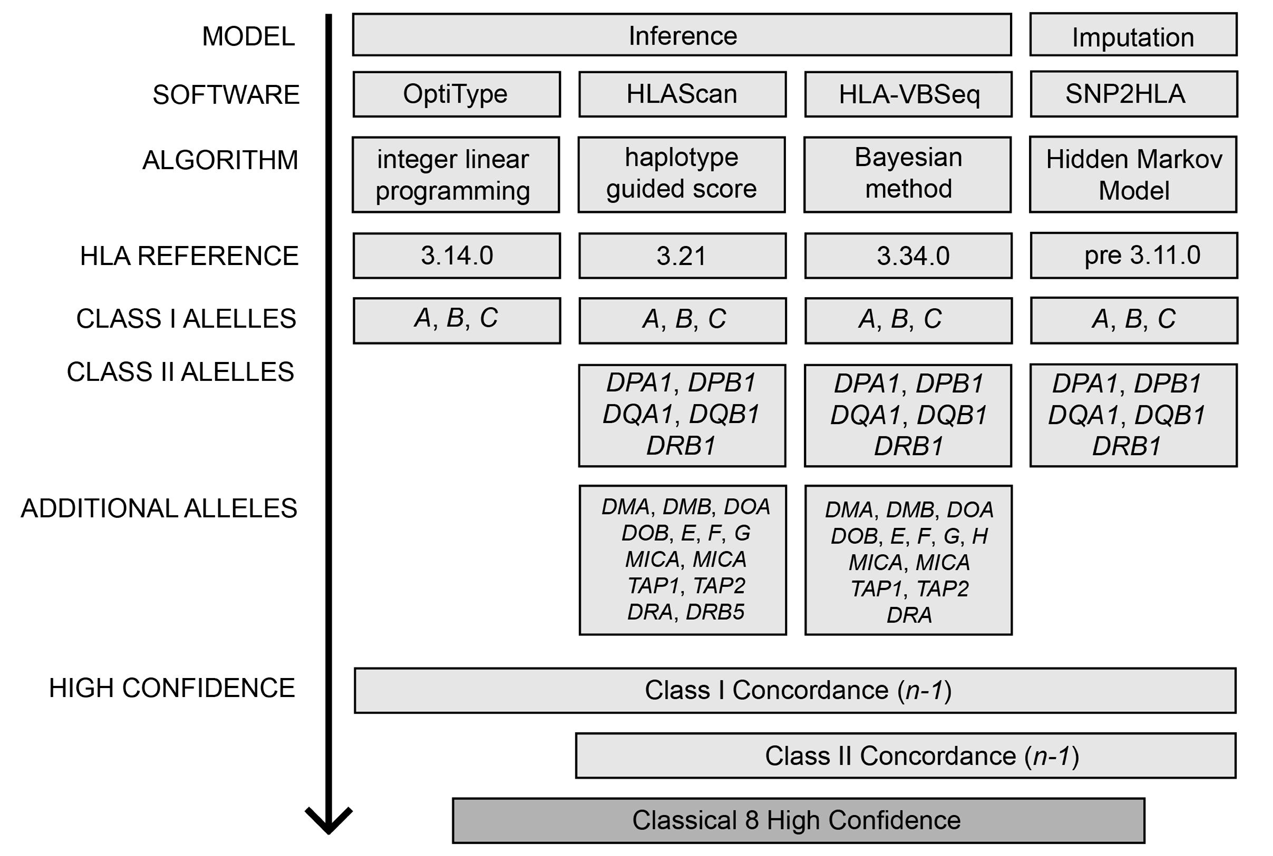
Pipeline for preliminary HLA allele typing and generation of a high confidence gene set. Four software solutions, representing two different models (imputation or inference) were used for typing. Each software utilised a separate algorithm and HLA reference to call a variable number of genes. An *n-1* concordance rule was used to create the high confidence set for the classical 8 genes (*HLA-A, -B, -C, -DPA1, -DPB1, -DQA1, -DQB1, -DRB1*).

SNP2HLA ^14^ utilizes the Hidden Markov model of Beagle ^19^ and the T1DGC reference panel of 5 225 Europeans ^14^ to impute HLA alleles based on a pre 3.11.0 reference (version not specified). The result is 2-field allele information for the classical 8 genes. The default settings, 10 iterations and window size of 1 000 markers, were implemented.

OptiType ^15^ views HLA typing as an optimization problem and uses an integer linear program to estimate which allele explains the largest number of reads. This software implements a custom-made IMGT/HLA v3.14.0 reference, where nucleotide sequences have been complemented with intronic information from the closest neighbour of the allele. These genomic-like sequences focus on exons 2 and 3 and allow for the calling of *HLA-A, -B* and *-C*, to a 2-field resolution.

HLA-VBSeq ^16^ uses a variational Bayesian approach to remap reads to a user defined IMGT/HLA genomic reference, we selected v3.34.0. Default settings were used, however the recommended allele coverage threshold (>20% of mean coverage) was relaxed to 10% in order to increase the number of alleles reported (reduced from 30% to 0.5% NA genotypes). Coverage was calculated for the 21 genes available in HLA-VBSeq using the longest transcript and Picard v1.92 *HS-metrics* (http://broadinstitute.github.io/picard/). HLA-VBSeq typed at up to a 4-field resolution.

HLAscan ^17^ realigns reads to a reference consisting of the nucleotide sequences from IMGT/HLA v3.21.0. It relies on a score function that ranks alleles based on the number of unique reads mapping to each, including a gap penalty. Alleles are reported up to a 3-field resolution, and calls are based on exon 2 and 3 for class I genes, and exon 2 for class II genes. Default settings were used; score cut-off 50, constant using ScoreFunc 20, for the 21 available genes.

### Creation of a high confidence call set

Each software has its own inherent biases, such as IMGT/HLA reference version, reference region considered, and the algorithm used. To reduce the impact of these, a high confidence HLA allele set was generated by merging results based on an *n-1* software concordance (Figure 1). An individual was classed as “typed” if the allelic pairs for three out of four software matched for class I, or two out of three for class II. Downstream population allele frequencies were calculated from SweHLA.

### Phasing of HLA haplotype blocks

SweHLA alleles were used as input to estimate haplotype blocks across the classical 8 genes with PHASE v2.1.1 ^20^. The -MS model ^21^ was run over 10 000 iterations using a thinning interval of 5 and a burn-in of 100. The model was run 10 times using different seeds for each. In order to maintain phasing power but reduce computational time, the eight genes were divided into three blocks based on known recombination hotspots (1: *HLA-C* and *-B*, 2: *HLA-DRB1, - DQA1* and *-DQB1*, and 3: *HLA-DPA1* and *-DPB1)* ^22^. In this way the maximum number of samples per block could also be considered. Haplotypes from the three intermediate blocks were combined in the following sequence, *HLA-A* with block 1, followed by block 2 and 3.

### Benchmarking the HLA typing accuracy and population frequency

Across software comparisons were performed to investigate the impact of software choice on the ability to call HLA alleles. SweHLA was assigned as the truth set and a concordance rate per allele calculated for each software. Concordance rate was defined by counting the number of times an allele was correctly called, divided by the total number of SweHLA calls for the same allele.

Within and across population comparisons were also conducted. We estimated allele calling accuracy by comparing SweHLA allele frequencies to an independent lab typed Swedish population ^9,10^. The lab typed set consisted of 252 unrelated individuals at 2-field resolution for *HLA-A, -B, -DQA1, -DQB1* and *-DRB1*. Correlation (r^2^) was calculated with *cor()* in the R v3.4 environment ^23^. To place SweHLA results in the context of Europe, gene and haplotype frequencies were compared to those of a recently published SNP2HLA imputed British population (5 544 individuals) ^11^.

## Results

### Characterisation of MHC region

The ability to call HLA alleles across the MHC is directly related to the quantity and quality of the reads mapped and variants called. Given the variability of coverage across this four Mb region (average 46.8 x; range 7.5-90.5 x; Figure 2A), we examined the repeat and gene content of the one kb bins sitting at the extremes of the distribution (coverage <20 x or >70 x). As expected, these regions predominantly contained repeat elements (61% were L1 LINEs, Alu SINEs and ERV1 LTRs), however we did note exons 1, 3 and 6 of *HLA-DRB1* (NM_002124.3) were covered with >70 x. These coverage extremes illustrate the inherent problems of mapping short-read data uniquely to repeats or across genes and paralogues.

**Figure 2.**
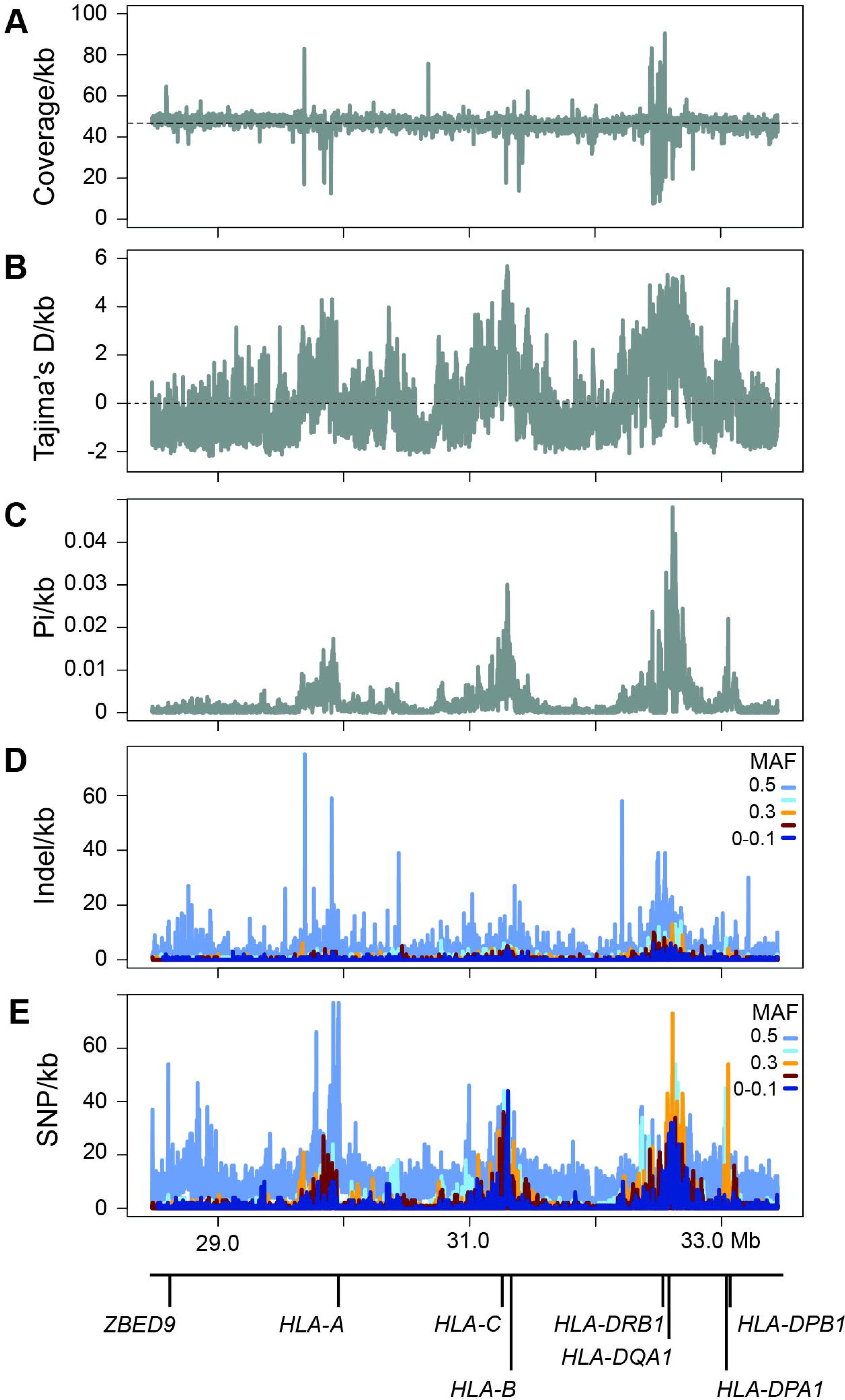
Characterisation of read depth and variation in 1-kb bins across the MHC. (A) Coverage peaks and troughs are illustrated in relation to the average across the region (46.8 x, dotted line). Metrics of genetic diversity, Tajima’s D (B) and Pi (C) are plotted for the same bins, as are density values for indels (D) and SNPs (E), although the latter are further dived into minor allele frequency (MAF) bins.

We used the metrics of Tajima’s D and Pi, in combination with variation density, to examine the patterns of selection and diversity across the MHC (Figure 2B-E). The three main peaks in each panel are centred over the class I (e.g. 29.9 Mb near *HLA-A* and 31.2 Mb near *HLA-C*) and class II genes (e.g. 32.6 Mb near *HLA-DQA1*), likely reflecting the selection pressure on these key immune gene classes (as Tajima’s D >3, Figure 2B). The strongest region of nucleotide diversity spanned the class II genes, with the apex including the 3’UTR of *HLA-DQA1* (NM_002122.3, Pi=0.048, 137 SNPs; Figure 2C). In contrast to both class I regions, the 32.6 Mb section contains the highest density of SNPs in the 0.2-0.3 and 0.4-0.5 MAF bins (orange and light blue respectively, Figure 2E). We further dissected the 87,637 variable positions in both the indel and SNP bins to characterise which fraction represented known (dbSNP v147) or novel variation (Supplementary Table S1). In each case, the majority of novel variation was found in the lowest MAF bin (MAF<0.1; Supplementary Table S1, 69.0% of novel indels and 58.3% of novel SNP; Figure 2D and E, dark blue band). For indels the fraction of novel variants per bin remained fairly steady (30-40%), however this value dropped markedly for SNPs (1.5-3.0%), and noticeably, only singletons not found in ExAC ^24^ were located in exons 2 or 3 of the classical 8 genes (data not shown).

### HLA alleles from four software and high confidence calls

Each of the software programs applied demonstrated a high per gene HLA typing rate, ranging between 98.1-100% (Supplementary Table 2). Per software the most difficult gene to call was *HLA-DPA1* (938 samples called by HLAscan, Supplementary Table 2), while for SweHLA it was *HLA-DQA1* (824 samples typed, Table 1). The overall genotyping rate dropped slightly for SweHLA (93.7%), however this was due to cross software mismatches and not a single individual’s inability to be typed. Of the small fraction of SweHLA alleles that were called as NA (1 006/16 000 alleles), most (*n*=863) could be resolved if typing was relaxed to the serological antigen 1-field level.

**Table 1.** Summary information for the high confidence set, SweHLA.

|  |  | Samples<sup>1</sup> | Alleles<sup>2</sup> | Homozygosity (%) | Coverage<sup>3</sup> |
| --- | --- | --- | --- | --- | --- |
| *HLA-* | *A* | 981 | 28 | 17.7 | 37.7 |
|  | *B* | 971 | 36 | 9.8 | 35.5 |
|  | *C* | 976 | 23 | 12.1 | 35.1 |
|  | *DQA1* | 824 | 15 | 15.0 | 35.6 |
|  | *DQB1* | 988 | 16 | 11.9 | 34.0 |
|  | *DRB1* | 901 | 33 | 11.1 | 32.3 |
|  | *DPA1* | 982 | 4 | 78.7 | 37.0 |
|  | *DPB1* | 875 | 20 | 28.9 | 36.8 |
| Gene set | Class I | 932 | NA | 2.6 | 36.1 |
|  | Classical 6 | 690 | NA | 1.2 | 35.0 |
|  | Classical 8 | 593 | NA | 0.5 | 35.5 |

<sup>1</sup>1000 samples were available for typing at each gene. <sup>2</sup>Total number of 2-field alleles typed at each gene. <sup>3</sup>Coverage was calculated as the part of the HLA-VBSeq threshold. NA, not applicable.

For the SweHLA class I gene set, 932 samples were called for all three genes, 60 samples for two genes and only four samples had one gene typed (Figure 3). A similar pattern emerged as we built up to the classical 8 through the classical 6 set. The poorer SweHLA calling at the *HLA-DQA1* locus impacted this set, for which 690 samples had genotypes for all six genes. However, a further 264 samples were typed at five genes, leaving only 45 samples with a minimum of three genes called (Figure 3). At the classical 8, 593 samples were typed at all genes, while 920 samples have high confidence calls at seven or more genes.

**Figure 3.**
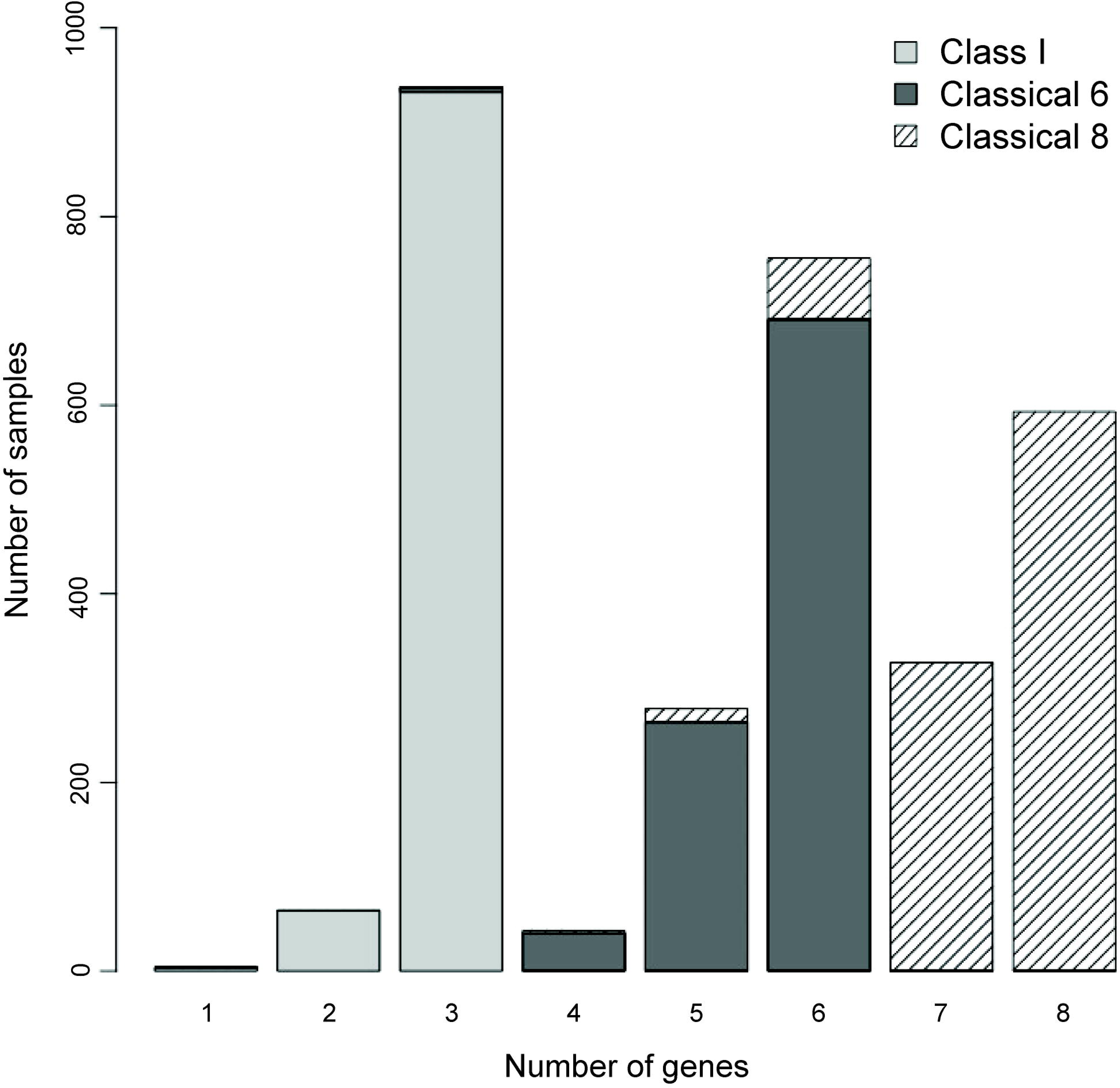
Distribution of samples successfully called in stages up to the creation of the complete SweHLA set. The majority of the 1 000 SweGen samples were called at all three class I genes (light grey, 3 genes *n*=932). Building up to the classical 6 genes, 690/1 000 samples were genotyped successfully at all loci (dark grey), which decreased slightly for the classical 8 (hashed, 8 genes *n*=593).

An average of 22 alleles were called for each of the eight genes investigated for SweHLA (range 4-36, Table 1). This was not related to the absolute number of alleles available per software, but rather to Swedish population diversity. For example, while between 298 and 9 854 2-field alleles were available across the software tested (SNP2HLA and HLAscan respectively, Supplementary Table 3), only 1.9% (class I) and 4.1% (class II) of the alleles were common across all programs (Supplementary Figure S1A-B). However, an intersection of the alleles called (Supplementary Table 3), revealed that for class I, 32.5% of alleles were typed in all software (37.7% in *n-1* programs) and for class II this fraction was 55.7% (72,1% for *n-1* programs, Supplementary Figure S1C-D).

We recorded population level frequencies for each gene and software combination (Supplementary Table S4) and noted that shared allele availability did not always translate to shared allele frequency. For example in class I genes, small frequency fluctuations were observed across data sets for *HLA-A* (*A*26:01* ranged between 1.8-2.5%; 2.1% SweHLA, Figure 4A), while larger discrepancies were noted for *HLA-B* (*B*27:05:* 4.8-8.0%; 7.6% in SweHLA, Supplementary Figure S2A). In class II, the variations were even larger and occurred more frequently. For *HLA-DRB1*, the most common allele *HLA-DRB1*15:01* (16.1%, Figure 4C), spanned a software range of 6.1-17.7%.

**Figure 4.**
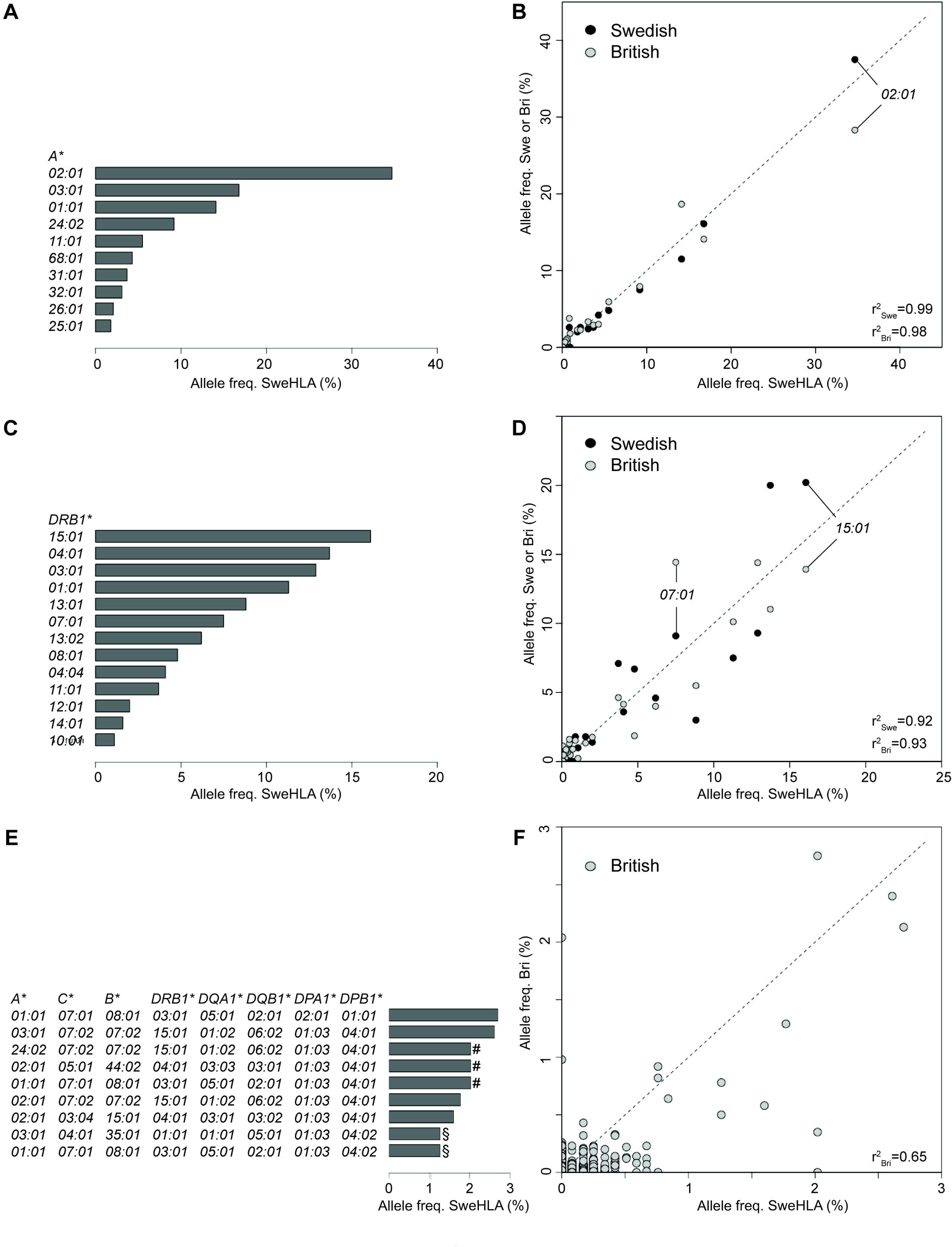
SweHLA allele frequency distribution and correlation (r^2^) to a separate lab typed Swedish (Swe) population and an imputed British (Bri) set. (A) *HLA-A*02:01* was the most frequent allele found in SweHLA (34.7%), as well as in (B) Swe and Bri, although at lower frequencies. (C) The frequency pattern in H*LA-DRB1* was mixed, with the sixth ranked *07:01* SweHLA allele the most frequent observed in (D) Bri, and ranked fourth in Swe. (E) All classical 8 allele SweHLA haplotypes were observed with a population frequency of <3%. With (#) and (§) indicating haplotypes with the same allele frequency. (F) At the single gene allele level, Bri and SweHLA were quite similar (r2>0.93 (B,D)), across haplotypes, there were haplotypes observed at >2% frequency in one population but absent from the other. SweHLA allele frequencies above 1% are plotted in bar graphs. Lab typed Swedish (Swe), *n* = 252, black circles; imputed British (Bri), *n* = 5 544, grey circles).

In order to explore discrepancies more thoroughly, we plotted the concordance rate per allele against the frequency per allele for each gene and software (Supplementary Figure S3). In general, SweHLA alleles observed at a frequency greater than 5% showed concordance above 90%. There were a few notable exceptions; *HLA-B*27:05* and *-DRB1*15:01* as mentioned previously, as well as *HLA-C*05:01*, two *-DPB1* and *-DQA1* alleles (Supplementary Figure S3B-E and H). In each case, SweHLA allele frequency was 7.6% or above, with a concordance rate below 80%. We noted previously that *HLA-DQA1* had the lowest genotyping rate (824/1 000 samples, Table 1) and Supplementary Figure S4E illustrates that this problem is in large part due to missing reference data; seven alleles with a population allele frequency ranging 0.06-6.00% were not present in the SNP2HLA reference (dark blue line). Combined, these alleles represent 11% of *HLA-DQA1* diversity in SweHLA.

Given that SNP2HLA is an imputation software, we investigated if the original alignment of reads to the reference could have affected SNP availability for this process. This may indeed have been the case. There are eight curated European HLA haplotypes available for alignment, with hg19 incorporating the PGF haplotype ^25^. This is important, as within exon 2 of *HLA-DQA1* there is a stretch of ∼100 nucleotides, common to COX and QBL, but lacking in the other five haplotypes including PGF. Mapping to hg19 results in the soft clipping of reads in this region and a dramatic drop in coverage (Supplementary Figure S4). The latter can affect allele calling in both homo- and heterozygotes, illustrated clearly when uncalled samples were aligned to either PGF, or alternate haplotypes (Supplementary Figure S4).

### Allele and haplotype correlations across populations

The SweHLA population frequencies for *HLA-A, -B, -C, -DQA1, -DQB1* and *-DRB1* were highly correlated with those of an independent lab typed Swedish population (r^2^ spans between 0.87-0.99 for *HLA-DQA1* to *–A*; black circles, Figure 4B, D and Supplementary figure S5A, C and E). There was no evidence that the number of alleles typed influence the correlation. *HLA-DQA1* has 15 alleles and *-DQB1 16,* yet r^2^ is 0.87 and 0.93 respectively. High levels of genetic homogeneity across HLA has been reported for Europe, with the diversity estimated to be as low as ∼5% ^26,27^. It was therefore not surprising that the frequency comparison between SweHLA and >5,500 British samples gave only slightly lower correlations than those to a Swedish population (r^2^ spans 0.83-0.98 for *HLA-DQA1* to *–A*; grey circles, Figure 4B, D and Supplementary Figure S5).

Phased MHC blocks can be used in multiple downstream analyses, including the imputation of missing allele calls, creating population reference graphs, the investigation of allele group interactions and to dissect disease causing mechanisms ^25,28,29^. In our phasing of the classical 8 gene haplotypes, only samples with complete allele typing per intermediate block were included. For example, block 1 (*HLA-C* and *-B*) utilised the results of 948 samples, whereas block 2 (*HLA-DRB1, -DQA1* and *–DQB1*) was reduced to 733 samples (Supplementary Figure S6A-C). At the resolution of the classical 6 genes (S5D Figure), it was revealed that COX (7%) and PGF (4%) were the most common haplotypes in SweHLA. This result could be further teased apart at the classical 8 haplotype level (Figure 4E), with PGF still among the most frequent at 2.6% of the total, however VAVY which shares the classical 6 haplotype with COX, became the most common haplotype (2.7%) ^29^.

The maximum allele frequency for the classical 6 haplotype was below 5% and for the classical 8 it was reduced to below 3% (Figure 4E, Supplementary Figure S6D). This was in keeping with the frequencies reported for the British population ^11^. Comparing across these populations, we can see the effect of recombination to shuffle common alleles to create rare haplotypes (classical 8 haplotype r^2^=0.648, Figure 4F), however COX and PGF were also the most frequent 6 gene haplotypes in the British population, with VAVY ranked third in the 8 gene haplotype ^11^.

### Significant differences found in homozygosity rate

We explored the homozygosity of SweHLA (Table 1) in comparison to other European derived populations. Per gene, SweHLA was compared to European Americans (*HLA-A, –B -C, -DQA1*, and *-DRB1)* ^30^ and for the classical 6 and 8 haplotypes SweHLA compared with same the British population as before ^11^. Both *HLA-A* and *–B* were significantly more homozygote than European Americans (population proportional Z-score; 17.7% vs 15.2%, *p-value*=0.044 and 9.8% vs 7.0%, *p-value*=0.002, respectively). For the classical 6 and 8 haplotypes, no significant differences to the British cohort were observed (*p-values*: 0.44 and 0.31).

## Discussion

Drawn from the 1 000 genomes of SweGen, SweHLA represents a high confidence bio-resource that provides a snapshot of Sweden’s MHC diversity. Data is reported at a clinically relevant resolution (2-field) and through the application of an *n-1* software concordance approach (Figure 1), is expected to have minimal allele bias. Results for the classical 8 genes are available at both the allele and haplotype level, and so SweHLA could be used to estimate HLA diversity within this population, or to tease apart the patterns of linkage disequilibrium surrounding these genes. As SweHLA is drawn from SweGen, and therefore also the Swedish Twin Registry, the resource’s value likely lies as an added control resource for the genetic dissection of disease linked to HLA genes. Access to raw genotyping or phenotypic data (sex, age, cohort) can be requested from each dataset mentioned following an individual review process.

SweHLA is a consensus resource; for allele calls to be reported, three out of four software matches were required for class I genes, relaxed to two out of three for class II genes. The absolute number of HLA typing programs considered was arbitrary, but reflected a range of factors that end users of all software should take into account, i) not all programs are developed to call the same gene set, ii) the IMGT/HLA allele reference employed may be fixed or dated, iii) algorithms differ between software.

Point one can be overcome through the selection of software to suit a specific need; although there are a lack of solutions developed to call outside class I, let alone the classical 8 genes set or those in the extended MHC region. Points two and three are perhaps the most clinically relevant, and reinforce the need for using multiple programs. Between 298 and 9 854 2-field reference alleles were available for the software programs we used, however only a small fraction of these were common to all (1.9% class I and 4.1% class II). While this fraction increased in our population after typing (32.5% class I and 55.7% class II), the choice of IMGT/HLA reference could lead to incorrect assumptions. For example, *HLA-DQA1*03:03* has a population frequency of 6% in SweHLA, but was not reported in the British population we used for comparison (Supplementary Figure S5C). This was not a reflection of diversity, rather the fact that *HLA-DQA1*03:03* is not available in the software used to analyse that dataset (Supplementary Figure S3). The flip side of this is when reference alleles are available, but not called due to the software’s algorithm. Here we use the example of *HLA-DRB1*16:01*, an allele previously associated with immune response in multiple sclerosis ^31^. In our hands, *HLA-DRB1*16:01* was called at a frequency of 0%, 0.3% and 10.7% depending on software choice (Figure 4E). It may therefore appear that a locus is not replicated, rather than mis-estimated.

These concerns were not limited to class II alleles. For *HLA-B*27:05*, an allele with high association to several diseases, including ankylosing spondylitis ^11,32^, we recorded allele frequency between 4.8 and 8.0%. Troublingly, one of the software had a concordance rate below 80% when compared with SweHLA (Supplementary Figure S3B). Others have noted that the *HLA-B*27* serotype has a higher frequency in the Nordic countries compared with other regions (>10% of *HLA-B* diversity ^9,33^), and so depending on population, this allele may appear rare (<5%) when in fact that is not the true case. These are not isolated examples. We noted 18 alleles with a delta allele frequency between any two HLA programs of more than 2%; 15 of these had reported association to at least one disease (data not shown).

The problem of variant calling from reads aligned to regions of high genetic diversity, high repeat content or containing paralogous genes is not new ^34,35^. A clear bias toward calling reference alleles was noted when HLA SNPs genotyped from 1000G (phase I) short read NGS were compared to those generated for the same individuals via Sanger sequencing ^36^. This trend was found in four of the five HLA genes examined, *HLA-A, -B, -DQB1* and -*C*, but not *-DRB1*^36^. It is here that population reference graphs ^29,37^ or alignment to the most similar MHC reference could aid variation discovery. We tested this at the *HLA-DQA1* locus with a subset of samples, typed or missing from SweHLA. First, we used the surrounding HLA classical 8 alleles to determine the most similar GRCh37 alternative haplotypes ^29^, and then aligned raw SweGen reads to those references and compared observable allelic variation. Supplementary Figure S4 illustrates how the problem of soft clipping in exon 2 can be resolved for homozygotes (e.g. SweGen_A) and some heterozygotes (e.g. SweGen_B), but the problem is more challenging for heterozygotes for which the alternate extended haplotype is not yet available (e.g SweGen_C). To overcome these issues, the community is developing software using HLA population graphs as the reference. These aim not only to improve HLA inference, but also identify novel alleles (e.g. HLA*PRG ^38^ and Kourami ^39^). However these provide G-group resolution, clustering alleles with identical sequence at the peptide biding domain.

While it was not the aim of this project to identify novel HLA alleles, we nonetheless examined the genetic diversity across the MHC for this population (Figure 2). Our results matched expectation, with the highest levels of Tajima’s D over the class I and II genes, and with more nucleotide diversity observed at class II genes compared to class I ^40,41^. The majority of SNP and indel variation fell into the 0-0.1 minor allele frequency bins. While a proportion of this will be true variation, as was noted above and by others, when the short read sequences are aligned to a more similar reference, a fraction of this variation will be revealed to be mapping errors ^42^.

With this work we have added to the growing set of HLA population resources now available for biomedicine. Whether the goal is to assess allele prevalence, dissect haplotype structure or develop a panel of additional control samples, the 1 000 genomes sourced to build SweHLA will be extremely valuable. Here the development of a *n-1* high concordance HLA set cleanly illustrates the need to apply more than one program to the problem of calling MHC alleles from short read data sets.

## Acknowledgements

We thank Mats Pettersson from Uppsala University for discussions around methodological biases. Computational resources were provided by the Swedish National Infrastructure for Computing (SNIC) at Uppsala Multidisciplinary Center for Advanced Computational Science (UPPMAX) under project sens2016003.

