## Supplementary Information for "SweHLA: the high confidence HLA typing bio-resource drawn from 1 000 Swedish genomes"

**Supplementary figures and tables**

Supplementary table S1. Distribution of known and novel variation across the MHC.

Supplementary table S2. Number of samples that were HLA typed by each software

Supplementary table S3. Summary of available and typed 2-field resolution reference alleles for each software.

Supplementary figure S1. Intersection of available and typed 2-field resolution reference alleles for each software

Supplementary figure S2. SweHLA allele frequencies for (A) *HLA-B*, (B) *HLA-C*, (C) *HLA-DPA1*, (D) *HLA-DPB1*, (E) *HLA-DQA1* and (F) *HLA-DQB1*.

Supplementary figure S3. Illustration of the relationship between allele frequency called per software and the high confidence SweHLA set.

Supplementary figure S4. Soft clipping is resolved when a sample is aligned to the most similar alternate HLA haplotypes.

Supplementary figure S5. Between population correlations for six of the classical 8 genes.

Supplementary figure S6. SweHLA haplotype frequencies at each of the three blocks used to construct the classical 8 gene haplotype.

Supplementary table S4. Population allele frequencies from each software as well as SweHLA

Supplementary Table S1. Distribution of known and novel variation across the MHC.

| Variant type | Bin[1] | rs[2] | non-rs[2] | % novel |
|---|---|---|---|---|
| Indel | 0.0 - 0.1 | 3720 | 7176 | 65.6 |
| | 0.1 - 0.2 | 849 | 578 | 40.5 |
| | 0.2 - 0.3 | 634 | 383 | 37.7 |
| | 0.3 - 0.4 | 424 | 241 | 36.2 |
| | 0.4 - 0.5 | 344 | 156 | 31.2 |
| | total | 5971 | 8534 | 58.8 |
| SNP | 0.0 - 0.1 | 29038 | 18804 | 39.3 |
| | 0.1 - 0.2 | 8645 | 202 | 2.3 |
| | 0.2 - 0.3 | 6879 | 211 | 3.0 |
| | 0.3 - 0.4 | 4982 | 150 | 2.9 |
| | 0.4 - 0.5 | 4158 | 63 | 1.5 |
| | total | 53702 | 19430 | 26.6 |

[1]Bins represent minor allele frequency (MAF) steps of 0.1 or the total content of MHC region. [2]rs designations are from dbSNP v147 as per SweGen.[1]

Supplementary Table S2. Number of samples that were HLA typed by each software

| Gene or gene set | HLAscan | HLA-VBSeq | SNP2HLA | OptiType |
|---|---|---|---|---|
| *A* | 1000 | 992 | 999 | 1000 |
| *B* | 1000 | 988 | 1000 | 1000 |
| *C* | 1000 | 986 | 1000 | 1000 |
| *DQA1* | 993 | 1000 | 998 | NA |
| *DQB1* | 987 | 1000 | 1000 | NA |
| *DRB1* | 971 | 999 | 1000 | NA |
| *DPA1* | 938 | 1000 | 1000 | NA |
| *DPB1* | 959 | 1000 | 1000 | NA |
| Class I MHC genes | 1000 | 968 | 999 | 1000 |
| Classical 6 | 951 | 967 | 994 | NA |
| Classical 8 | 854 | 967 | 997 | NA |

Maximum number of samples available, 1000. NA, not applicable.

Supplementary Table S3. Summary of available and typed 2-field resolution reference alleles for each software.

| | Available | | | | Typed | | | |
|---|---|---|---|---|---|---|---|---|
| Gene | HLAscan | HLA-VBSeq | SNP2HLA | OptiType | HLAscan | HLA-VBSeq | SNP2HLA | OptiType |
| *A* | 2382 | 947 | 50 | 1678 | 40 | 62 | 32 | 28 |
| *B* | 3048 | 1053 | 97 | 2273 | 52 | 48 | 42 | 46 |
| *C* | 2026 | 1077 | 33 | 1311 | 34 | 64 | 22 | 25 |
| *DPA1* | 20 | 13 | 7 | NA | 4 | 4 | 5 | NA |
| *DPB1* | 462 | 127 | 34 | NA | 22 | 30 | 21 | NA |
| *DQA1* | 33 | 26 | 8 | NA | 21 | 15 | 8 | NA |
| *DQB1* | 559 | 112 | 18 | NA | 17 | 21 | 16 | NA |
| *DRB1* | 1324 | 46 | 51 | NA | 37 | 29 | 35 | NA |

NA, not applicable.

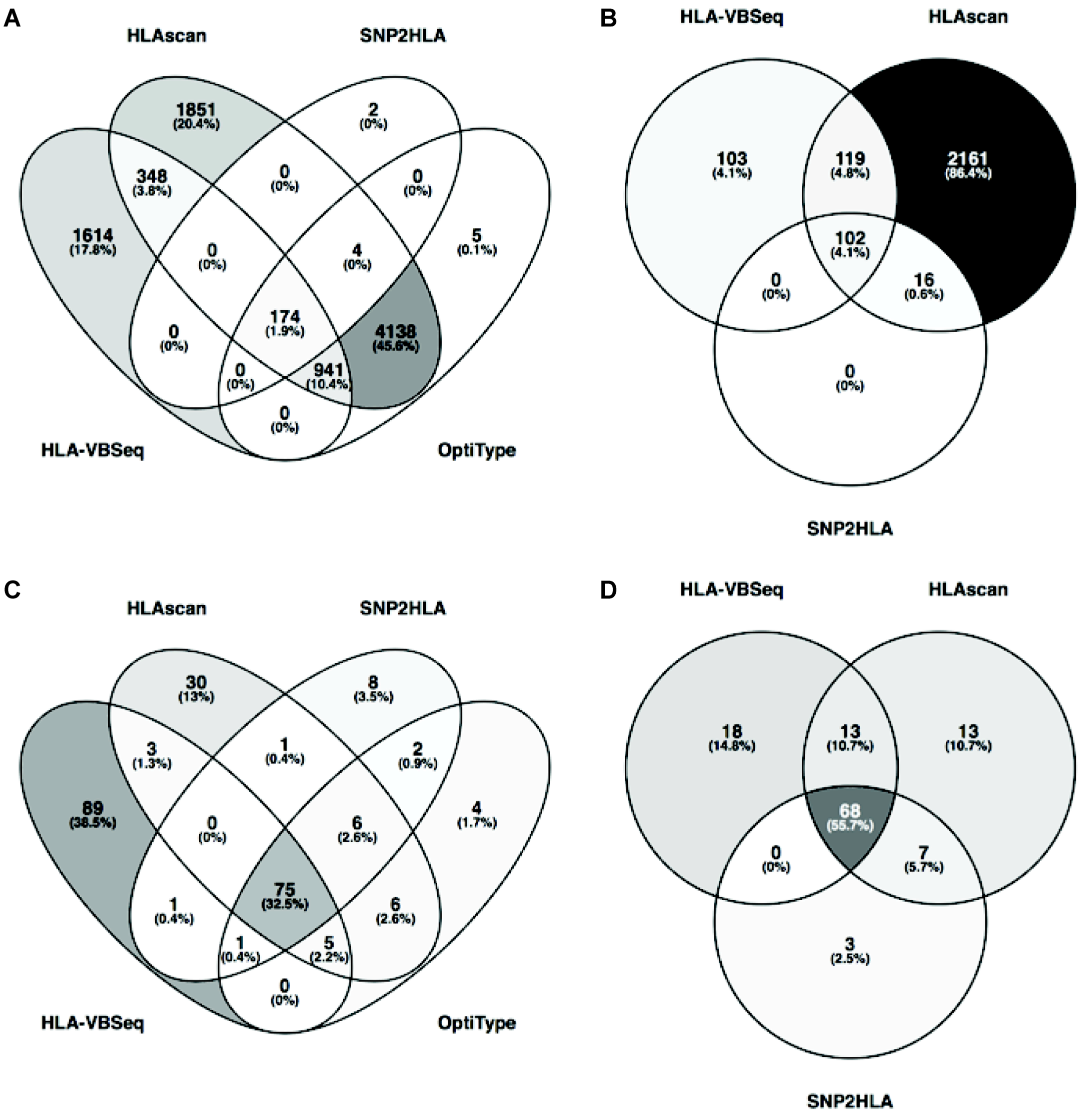


Supplementary Figure S1. Intersection of available; A) class 1 and B) class 2, and typed; C) class 1 and D) class 2 2-field resolution reference alleles for each software.

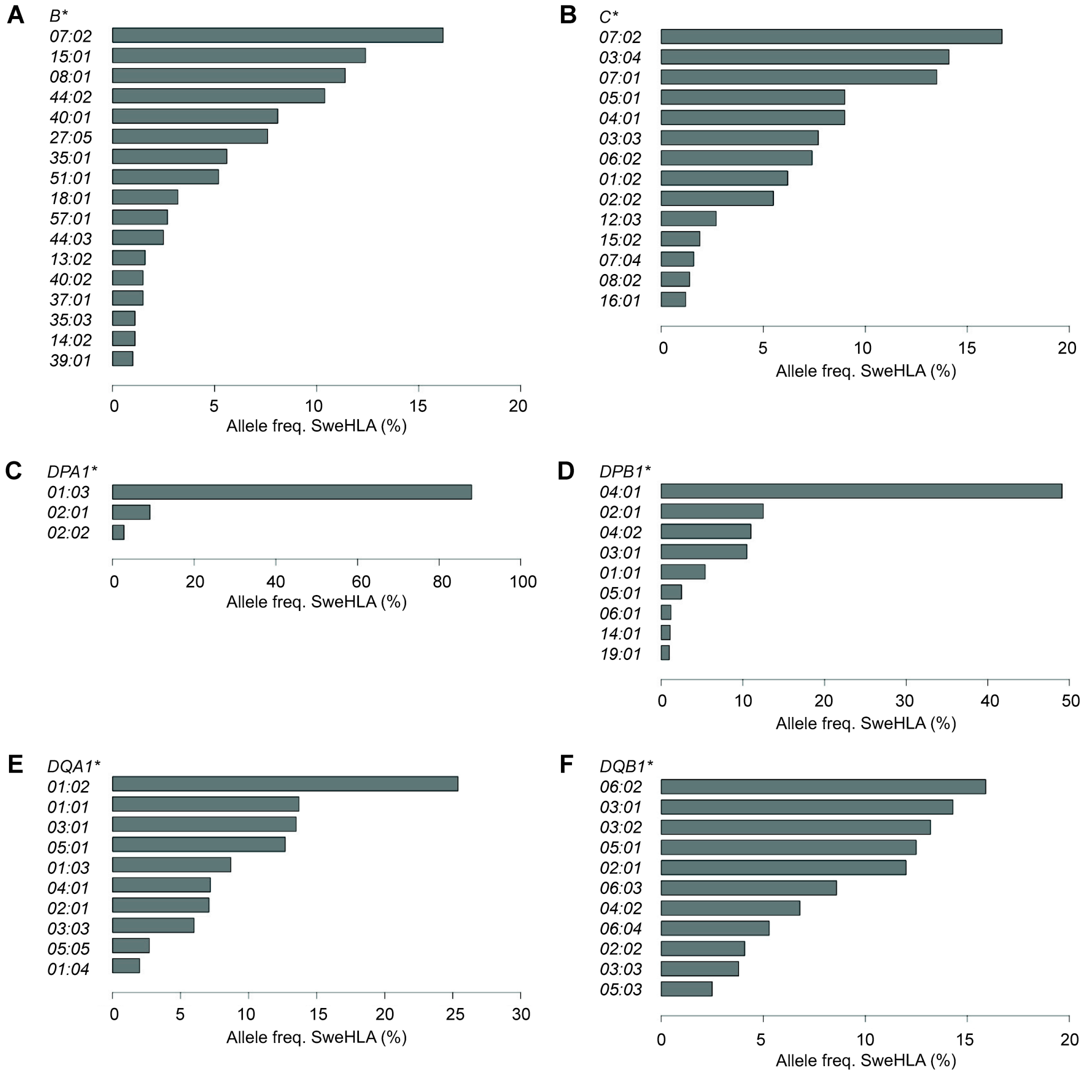


Supplementary Figure S2. SweHLA allele frequencies for (A) *HLA-B*, (B) *HLA-C*, (C) *HLA-DPA1*, (D) *HLA-DPB1*, (E) *HLA-DQA1* and (F) *HLA-DQB1*. Alleles with frequency greater than 1% are illustrated.

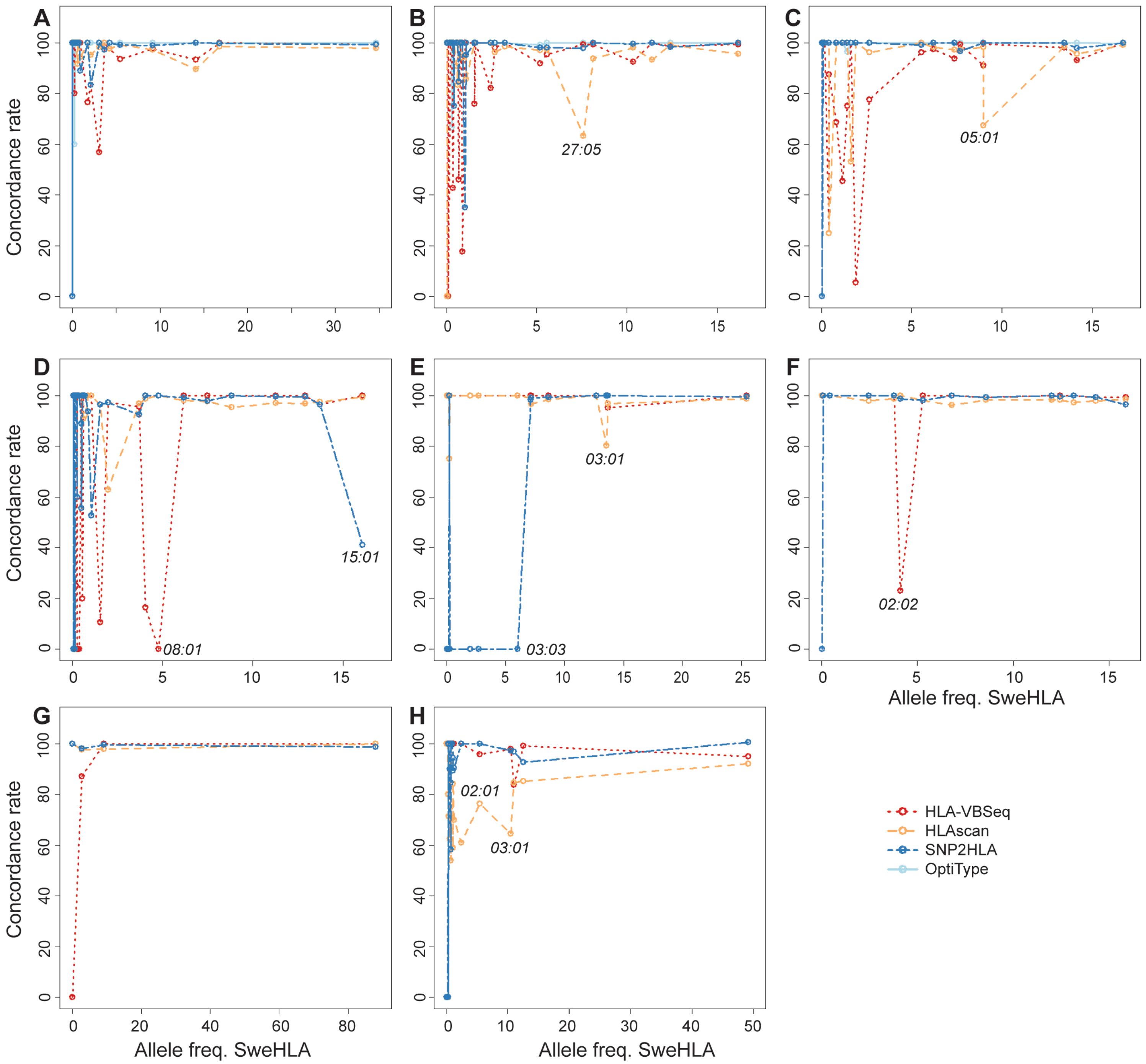


Supplementary Figure S3. Illustration of the relationship between allele frequency called per software and the high confidence SweHLA set. (A) *HLA-A*, (B) *HLA-B*, (C) *HLA-C*, (D) *HLA-DRB1*, (E) *HLA-DQA1,* (F) *HLA-DQB1*, (G) *HLA-DPA1* and (H) *HLA-DPB1*. Software used for genotyping were HLA-VBSeq (red), HLAscan (yellow), SNP2HLA (dark blue) and OptiType (light blue).

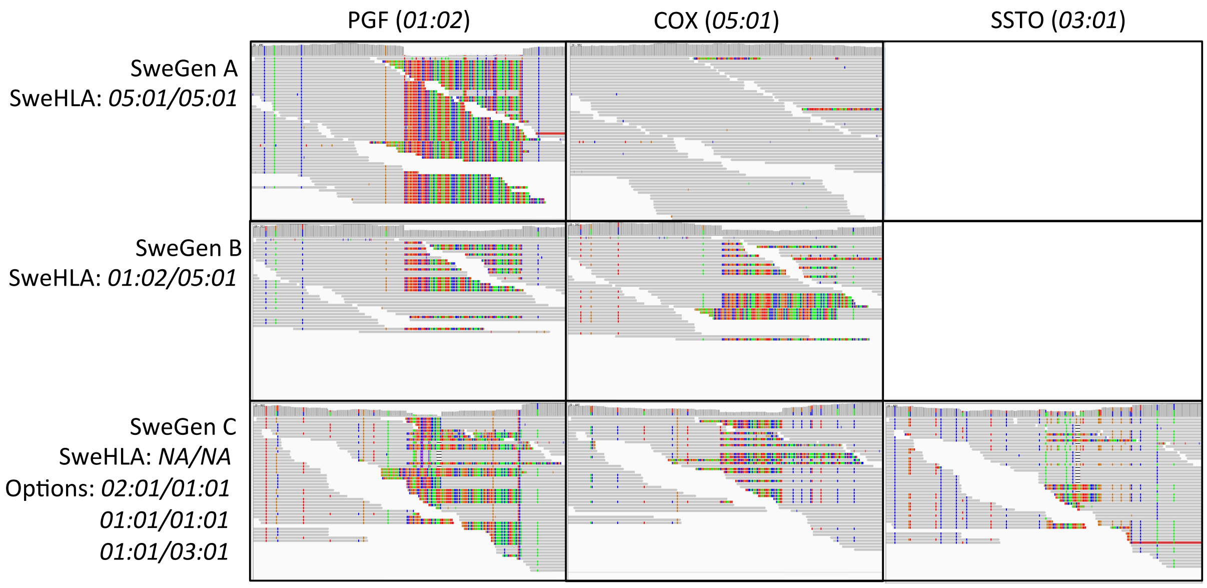


Supplementary Figure S4. Soft clipping is resolved when a sample is aligned to the most similar alternate HLA haplotypes. The zoomed region represents HLA-DQA1 exon 2 for PGF, COX and SSTO cell lines, with allele typing sourced from [2].

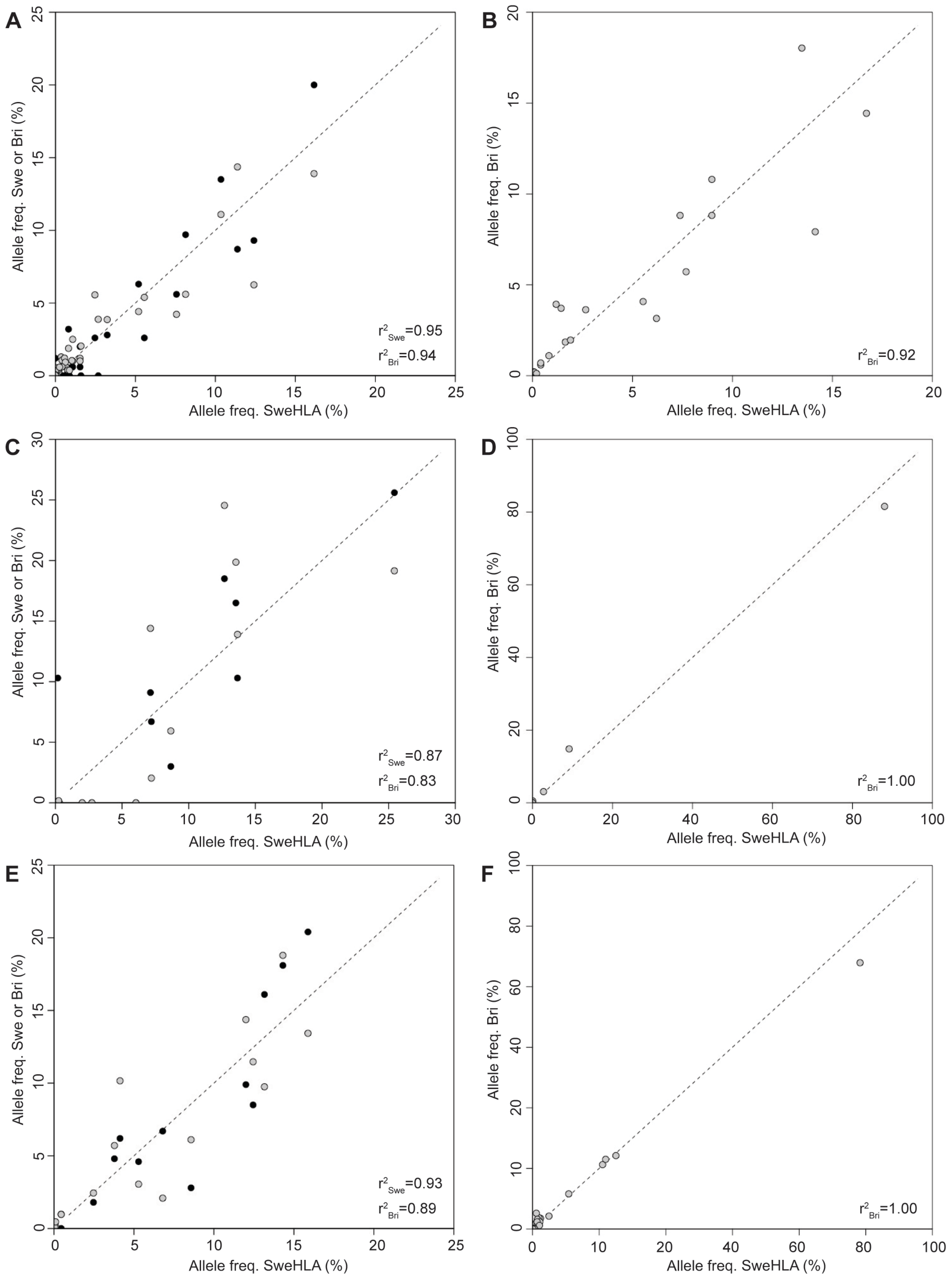


Supplementary Figure S5. Between population correlations for six of the classical 8 genes. Illustrated are SweHLA versus a lab typed Swedish cohort (Swe, black) and versus a British resource imputed with SNP2HLA (Bri, grey). (A) *HLA-B*, (B) *HLA-C*, (C) *HLA-DQA1*, (D) *HLA-DPA1*, (E) *HLA-DQB1,* (F) *HLA-DPB1*.

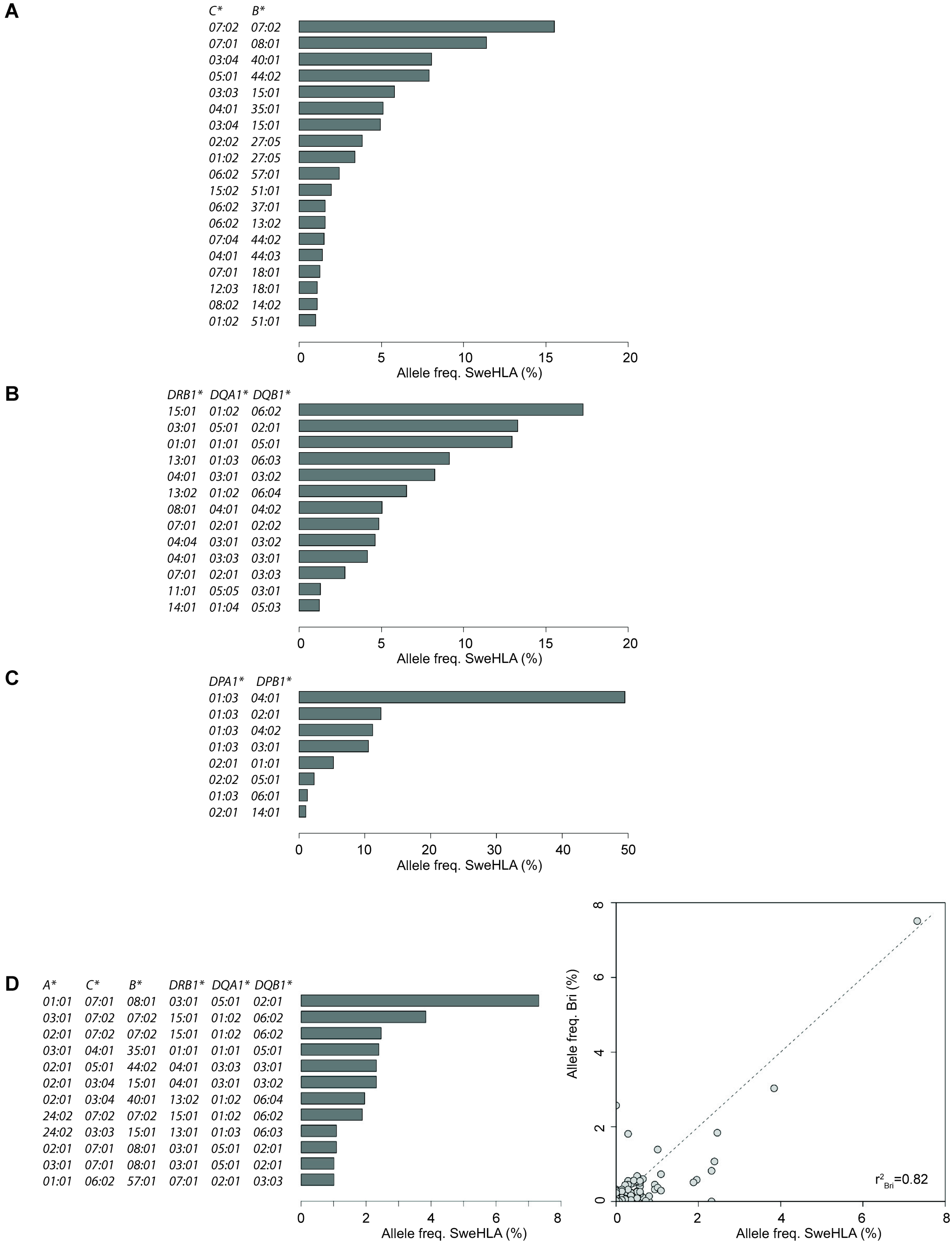


Supplementary Figure S6. SweHLA haplotype frequencies at each of the three blocks used to construct the classical 8 gene haplotype. (A) Block 1, *HLA-B*, *-C*, (B) *HLA-DRB1, -DQA1, -DQB1* and (C) *HLA-DPA1, -DPB1*. (D) The SweHLA haplotypes frequencies for the classical 6 genes are plotted, as is the correlation between these and the British resource imputed with SNP2HLA (Bri, grey). Frequencies above 1% are plotted in bar graphs.
